# Structural control energy of resting-state functional brain states reveals inefficient brain dynamics in psychosis vulnerability

**DOI:** 10.1101/703561

**Authors:** Daniela Zöller, Corrado Sandini, Marie Schaer, Stephan Eliez, Danielle S. Bassett, Dimitri Van De Ville

## Abstract

How the brain’s white-matter anatomy constrains brain activity is an open question that might give insights into the mechanisms that underlie mental disorders such as schizophrenia. Chromosome 22q11.2 deletion syndrome (22q11DS) is a neurodevelopmental disorder with an extremely high risk for psychosis providing a test case to study developmental aspects of schizophrenia. In this study, we used principles from network control theory to probe the implications of aberrant structural connectivity for the brain’s functional dynamics in 22q11DS. We retrieved brain states from resting-state functional magnetic resonance images of 78 patients with 22q11DS and 85 healthy controls. Then, we compared them in terms of persistence control energy; i.e., the control energy that would be required to persist in each of these states based on individual structural connectivity and a dynamic model. Persistence control energy was altered in a broad pattern of brain states including both energetically more demanding and less demanding brain states in 22q11DS. Further, we found a negative relationship between persistence control energy and resting-state activation time, which suggests that the brain reduces energy by spending less time in energetically demanding brain states. In patients with 22q11DS, this behavior was less pronounced, suggesting a dynamic inefficiency of brain function in the disease. In summary, our results provide initial insights into the dynamic implications of altered structural connectivity in 22q11DS, which might improve our understanding of the mechanisms underlying the disease.

## 1. Introduction

The brain is a complex dynamic system and brain function during rest and task can be described in terms of the dynamic activation and interaction of different *brain states*: sets of brain regions that are coherently activating and deactivating (Preti et al., 2017; Karahanoğlu and Van De Ville, 2017). More specifically, healthy brain function is characterized by continuous transitions between different cognitive states (as for example the internally oriented default mode network (DMN), or the attention-related fronto-parietal network (FPN)), and alterations in these dynamic state transitions could inform on disrupted brain function in mental diseases (Christoff et al., 2016). How the brain’s underlying structural backbone constrains and facilitates this dynamic behavior is an intensively studied question in the neuroscience community (Bassett and Sporns, 2017; Honey et al., 2009; Becker et al., 2018). The joint consideration of structural and functional properties is particularly promising to provide a better mechanistic explanation of the causes that underlie brain disorders such as schizophrenia (Braun et al., 2018). In recent years, approaches for the investigation of dynamic properties have proven to be particularly useful in probing brain function in health and disease (Preti et al., 2017; Karahanoğlu and Van De Ville, 2017; Van Den Heuvel and Fornito, 2014). Schizophrenia, in particular, is – as an extension of the well-accepted dysconnectivity hypothesis (Friston et al., 2016) – increasingly perceived as a disorder of broad alterations in large-scale brain state dynamics (Fornito et al., 2012; Du et al., 2016; Braun et al., 2016). As schizophrenia, and mental disorders in general, have a polygenetic basis in combination with strong social-environmental and developmental implications, they are likely also affecting a multitude of biological systems(Braun et al., 2018). Thus, a better insight on how the alterations in the brain’s structure may lead to aberrant dynamic activation would improve our understanding of this disease to ultimately improve clinical management and patient outcomes (Braun et al., 2018).

Network control theory provides a framework to address this question of structure-function relationship, and to analyze how the brain’s structural topology influences its dynamic function (Gu et al., 2015; Betzel et al., 2016; Gu et al., 2017; Kim et al., 2018; Karrer et al., 2019). While initial studies of the relationship between brain structure and function used cross-modal correlations (Honey et al., 2009), network control theory approaches aim to mechanistically describe how brain structure influences its function (Bassett and Sporns, 2017; Braun et al., 2018). In network control theory approaches, the brain is modeled as a graph defined by its structural connectivity, where nodes correspond to brain regions, and edges indicate structural connectivity strength between brain regions. The state of this system is defined by the neurophysiological activation of each brain region, and a (usually linear) dynamic model describing the dynamic transition between these functional brain states over time. Under the assumption that the brain’s state is controlled through an internal or external control input signal, it is then possible to quantify how the underlying structural architecture facilitates or constrains the system’s dynamic behavior. In other words, the measured brain state at one moment (in terms of hemodynamic activation) is modeled as the propagated activation during an earlier brain state and an added control input at the control regions. This assumption is the basis for many models of information propagation in the brain (Srivastava et al., 2020) and the validity of this view is supported by empirical evidence showing that regional fMRI activation levels during task can accurately be predicted by flow of activity between brain regions (Cole et al., 2016). Through this framework, network control theory can be used to investigate how much energy the brain would require (i.e., how high the control input would be) to steer itself into different cognitive states (through internal control inputs) (Gu et al., 2015; Betzel et al., 2016), or to identify the most promising targets for neurostimulation to control the brain’s state (through external control inputs) (Muldoon et al., 2016; Khambhati et al., 2019).

Of note, the network control theory allows to examine the control energy that is required for specific trajectories between a precisely defined initial state and a precisely defined target state. A simple intuitive example of such a state transition is the transition from activation of the DMN to activation of the FPN (Betzel et al., 2016; Gu et al., 2017; Kim et al., 2018). The initial and target states can either be defined by an atlas (Gu et al., 2017; Cui et al., 2020), or by data-driven brain states retrieved from functional magnetic resonance imaging (fMRI) (Braun et al., 2019; Cornblath et al., 2020). By defining identical initial state and target state, one can obtain the control energy that would be required to *persist* in any particular brain state (Cornblath et al., 2020). Despite it’s simplifying assumptions, the network control theory framework has the advantage that it allows to go beyond a purely correlative description of structure-function relationships and integrate functional and structural properties in a common model, without the use of more complex and computationally demanding generative models as for example (Kringelbach et al., 2020; Deco et al., 2018; Ghosh et al., 2008). A validation of the model was provided in a recent study that compared network control energy to brain state transition probabilities during rest and during an n-back working memory task (Cornblath et al., 2020). The authors of the study found a negative relationship between control energy and transition probability during rest, which supports that the linear diffusion of brain activity on the brain’s structural white matter connections indeed constrains brain state transitions at rest (Cornblath et al., 2020).

Studies in healthy and clinical populations have demonstrated that network controllability measures can provide characteristic profiles for different cognitive brain systems (Gu et al., 2015), change with development (Tang et al., 2017; Cui et al., 2020), are a reliable and heritable property of the structural connectome (Lee et al., 2020), and track individual profiles in cognitive functions (Lee et al., 2020), executive functions (Cui et al., 2020) and impulsivity (Cornblath et al., 2018). Additionally, controllability properties relate to different neurotransmitter profiles (Shine et al., 2019) and are promising to guide target selection for neurostimulation (Muldoon et al., 2016; Khambhati et al., 2019). Persistence control energy in healthy subjects was found to be significantly lower when computed for structural brain networks than for random null models preserving topology and spatial constraints, suggesting that the brain’s structural wiring promotes persistence in functional brain states at low energy cost (Cornblath et al., 2020). In patients with temporal lobe epilepsy, controllability was found to be reduced suggesting a lower range of functional dynamics (Bernhardt et al., 2019), and in patients with bipolar disorder fronto-limbic subnetworks with aberrant structural connectivity lead to altered controllability (Jeganathan et al., 2018). In patients with schizophrenia, the persistence control energy of a data-driven working memory brain state was found to be higher compared to healthy control subjects, which was interpreted as reduced stability of this brain state in patients (Braun et al., 2019). Further, this study also found evidence supporting that persistence control energy is modulated by dopamine D1 receptor expression, whereas dopamine D2 receptor expression was modulating control energy for brain state transitions.

When brain states are derived from fMRI, network control theory allows to directly compare the dynamic temporal properties of these brain states (measured during fMRI) with the persistence control energy that would be needed to engage in these same brain states based on structural connectivity (measured with diffusion weighted MRI) (Cornblath et al., 2020). In this way, it is possible to investigate if the predicted effect of aberrant structural brain architecture on brain dynamics (reflected by persistence control energy) is in line with the measured temporal activation during rest. In the present study, we use this framework in order to explore whether and how such an integration of network control theory with dynamic functional brain state analysis could inform on disrupted structure-function relationship relevant to the pathophysiology of psychiatric disorders. To this aim, we investigated individuals with 22q11DS, a neurodevelopmental disorder characterized by a 30-fold increased risk for developing schizophrenia (McDonald-McGinn et al., 2015). Due to the 30-40 % prevalence of schizophrenia by adulthood (Schneider et al., 2014), the disorder is considered a model for the investigation of developmental risk factors before the onset of full-blown schizophrenia (Insel, 2010; Bassett and Chow, 1999).

In 22q11DS, the white matter microstructure and connectivity has been extensively studied, mostly in terms of whole-brain or tract-based diffusivity properties (reviewed in Scariati et al., 2016). Affected whitematter bundles mostly include long-range frontal-frontal, frontal-occipital, and fronto-parietal connections (Scariati et al., 2016; Kikinis et al., 2016; Tylee et al., 2017; Olszewski et al., 2017; Roalf et al., 2017). Only a few studies have thus far examined the characteristics of structural whole-brain networks (Ottet et al., 2013b; Kikinis et al., 2013; Ottet et al., 2013a; Padula et al., 2017a; Váša et al., 2016; Zhan et al., 2018), also mostly reporting fronto-temporal, fronto-parietal (Ottet et al., 2013b,a; Zhan et al., 2018), and limbic dysconnectivity (Ottet et al., 2013a; Padula et al., 2015). From a topological perspective, Ottet et al. reported longer path lengths and disconnectivity of the brain’s hub regions, specifically, in the frontal lobes (Ottet et al., 2013b), and Váša et al. uncovered a ‘de-centralization’ in 22q11DS with a rerouting of shortest network paths to circumvent an affected core that included frontal, parietal, and subcortical regions (Váša et al., 2016). FMRI studies in 22q11DS have so far mostly focused on static properties. In the only three studies who looked at dynamic features of resting-state brain function, we found global reductions in variability of blood-oxygenation level dependent (BOLD) signals (Zöller et al., 2017b), and reduced BOLD variability in the dorsal anterior cingulate cortex (a central node of the salience network) in relationship to higher positive psychotic symptoms (Zöller et al., 2017a). In another study investigating dynamic properties of resting-state brain states, rather than voxel-wise BOLD variability, we found that in 22q11DS higher positive psychotic symptoms come with higher salience network activation and coupling, while higher levels of anxiety are related to higher activation of the amygdala and hippocampus (Zöller et al., 2019). In these studies, we demonstrated the implication of resting-state brain dynamics in psychopathology in 22q11DS. However, which effect the brain’s structural wiring has on these aberrant functional findings remains an open question.

Only two studies in 22q11DS to date have investigated structural and functional properties at the same time (Padula et al., 2015, 2017b) and none so far have attempted to examine the dynamic implications of an altered structural network architecture. Here, we bridge this gap by combining dynamic fMRI analysis with whole-brain tractography and principles from network control theory to investigate how the brain’s white matter connectivity may influence its dynamic behavior and how this relationship is affected in patients with 22q11DS. More specifically, we extracted brain states from resting-state fMRI scans similarly to our previous study focusing on brain state dynamics (Zöller et al., 2019). Then, we calculated the control energy that is required – based on the structural connectivity of the same subjects – to persist in these specific brain states (Braun et al., 2019; Cornblath et al., 2020). Our aim was to go beyond a pure correlative investigation of the relationship between structural and functional connectivity, and to probe the potential implications of aberrant brain structure on the brain’s dynamic activation by using a mechanistic model of structure-function relationships. We take advantage of the network control theory framework to directly compare the measured temporal properties of functional brain states during resting-state fMRI, with the predicted control energy to persist in this brain state based on the structural architecture. This persistence control energy is defined as the magnitude of the required internal control input signal as predicted by the dynamic model. Through this approach, we explore whether and how the network control modeling framework could inform about disrupted structure-function relationship in psychiatric disorders.

## 2. Materials and methods

### 2.1. Participants

FMRI analyses in this study were conducted on the identical dataset as Zöller et al. (2019), which included 78 patients with 22q11DS and 85 healthy controls (HCs), aged between 8 and 30 years. For structural connectivity analysis, we used diffusion MRI (dMRI) scans acquired from the same subjects. One patient with 22q11DS and 7 HCs had to be excluded from dMRI analyses because no dMRI scan was recorded for them. Table 1 shows demographic information of the remaining 77 patients with 22q11DS and 78 HCs for which both fMRI and dMRI scans were available.

**Table 1:** Participant demographics. N/A = not applicable.

|  | **HC** | **22q11DS** | **p-value** |
| --- | --- | --- | --- |
| Number of subjects (M/F) | 78 (32/46) | 77 (37/40) | 0.379 ($\chi^2$) |
| Age mean ± SD<br>(range) | 16.39 ± 5.52<br>(8.1-30.0) | 17.23 ± 5.39<br>(8.1-29.7) | 0.342 |
| Right handed* | 80.00% | 77.61% | 0.564 ($\chi^2$) |
| IQ** | 110.80 ± 13.75 | 70.32 ± 12.18 | <0.001 |
| **Number of subjects meeting criteria for**<br>**psychiatric diagnosis** | **N/A** | **42 (55%)** |  |
| Anxiety disorder | N/A | 9 |  |
| Attention deficit hyperactivity<br>disorder (ADHD) | N/A | 8 |  |
| Mood disorder | N/A | 5 |  |
| Schizophrenia or schizoaffective disorder | N/A | 3 |  |
| More than one psychiatric disorder | N/A | 17 |  |
| Anxiety & ADHD & Mood & Schizophrenia |  | 1 |  |
| Anxiety & ADHD & Mood |  | 1 |  |
| Anxiety & ADHD |  | 11 |  |
| Anxiety & Mood |  | 3 |  |
| Anxiety & Schizophrenia |  | 1 |  |
| **Number of subjects medicated** | **0** | **16** |  |
| Methylphenidate | 0 | 9 |  |
| Anitpsychotics | 0 | 2 |  |
| Anticonvulsants | 0 | 1 |  |
| Antidepressants | 0 | 1 |  |
| More than one class of medication | 0 | 3 |  |
| Antipsychotics & Anticonvulsants |  | 1 |  |
| Antipsychotics & Antidepressants |  | 1 |  |
| Methylphenidate & Antidepressants |  | 1 |  |

\* Handedness was measured using the Edinburgh laterality quotient, right handedness was defined by a score of more than 50. \*\* IQ was measured using the Wechsler Intelligence Scale for Children–III (Wechsler, 1991) for youth and the Wechsler Adult Intelligence Scale–III (Wechsler, 1997) for adults.

Prodromal positive and negative psychotic symptoms were assessed only in patients with 22q11DS by means of the structured interview for prodromal symptoms (SIPS; Miller et al., 2003).

### 2.2. Image acquisition

MRI scans were recorded at the Centre d’Imagerie BioMédicale (CIBM) in Geneva on a Siemens Trio (12-channel head coil) and a Siemens Prisma (20-channel head coil) 3 Tesla scanner. Supplementary Table S1 contains the number of scans that were recorded before and after the update to the Prisma scanner for each image modality, respectively. There was no significant scanner-by-group interaction.

Anatomical images were acquired using a T1-weighted sequence with 192 coronal slices (volumetric resolution = 0.86×0.86×1.1 mm^3^, TR = 2500 ms, TE = 3 ms, flip angle = 8°, acquisition matrix = 256×256, field of view = 23.5 cm, slice thickness = 1.1 mm, phase encoding R>>L, no fat suppression).

FMRI scans were recorded with an 8 minute resting-state session using a T2*-weighted sequence at a TR of 2.4 s (200 frames, volumetric resolution = 1.84×1.84×3.2 mm^3^, TE=30 ms, flip angle = 85°, acquisition matrix 94×128, field of view 96×128, 38 axial slices, slice thickness = 3.2 mm, phase encoding A>>P, descending sequential ordering, GRAPPA acceleration mode with factor PE = 2). Subjects were instructed to fixate on a cross on the screen, let their minds wander, and not fall asleep.

DMRI scans were acquired in 30 directions using a single shell diffusion-weighted sequence (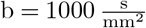, one b=0 image, volumetric resolution = 2×2×2 mm^3^, TR = 8300 ms to 8800 ms, TE = 84 ms, flip angle = 90° to 180°, acquisition matrix = 128×128, field of view = 25.6 cm, 64 axial slices, slice thickness = 2 mm, phase encoding A>>P, GRAPPA acceleration mode with factor PE = 2).

The cerebellum was not entirely captured in the resting-state scan for every individual and was thus excluded from the analysis.

### 2.3. fMRI processing

#### Preprocessing

We preprocessed fMRI scans identically as in Zöller et al. (2019) in Matlab R2017a using in-house code and functions of statistical parametric mapping (SPM12, http://www.fil.ion.ucl.ac.uk/spm/), Data Processing Assistant for Resting-State fMRI (DPARSF; Yan and Yufeng, 2010) and Individual Brain Atlases using Statistical Parametric Mapping (IBASPM; Aleman-Gomez et al., 2006). Briefly, preprocessing steps included functional realignment (registered to the mean) and spatial smoothing with a Gaussian kernel of 6 mm full width half maximum, co-registration of the structural T1-weighted image to the functional mean, segmentation of anatomical scans using the *Segmentation* algorithm in SPM12 (Ashburner and Friston, 2005), creation of a study-specific template using (DARTEL; Ashburner, 2007), exclusion of the first 5 functional frames, and regression of cerebrospinal fluid and white matter BOLD signals. Volumes with a framewise displacement (FD) larger than 0.5 mm were replaced with the spline interpolation of previous and following frames in order to ensure the constant sampling rate required for the retrieval of functional brain states using innovation-driven co-activation patterns (iCAPs, see following paragraph). After brain state extraction, motion frames were excluded for the computation of temporal characteristics (see below).

#### Extracting functional brain states: innovation-driven co-activation patterns

Following preprocessing, we extracted functional brain states from resting-state fMRI scans using the iCAPs method (Karahanoglu and Van De Ville, 2015; Zöller et al., 2018), for which code is openly available at https://c4science.ch/source/iCAPs. Briefly, steps to derive functional brain states (called “iCAPs”) and their temporal properties included the hemodynamically-informed deconvolution of fMRI timeseries using total activation (Karahanoglu et al., 2011, 2013; Farouj et al., 2017). Then, significant transients were determined with a two-step thresholding procedure following Karahanoglu and Van De Ville (2015) and Zöller et al. (2018) (temporal threshold: 5–95 %; spatial threshold: 5% of gray matter voxels). Functional brain states – the iCAPs – were then determined through temporal clustering on concatenated transient frames of all subjects. According to consensus clustering (Monti et al., 2003), the optimum number of clusters in the present study was *K* = 17. Finally, a time course was estimated for each iCAP using spatio-temporal transient-informed regression with soft assignment factor *ξ* = 1.3 (Zöller et al., 2018). For a more detailed description of the method, we refer the interested reader to (Zöller et al., 2019), where we used the identical approach on a largely overlapping sample.

#### Calculation of temporal activation duration

The temporal activation duration of each brain state (iCAP) was computed from it’s thresholded time course (z-score *> |*1*|*). We defined the activation duration as the percent of active timepoints (amplitude *> |*1*|*) with respect to the total non-motion scanning time.

### 2.4. dMRI processing

DMRI scans were processed using functions from the FSL library (v5.0.11; Jenkinson et al., 2012) and from the MRtrix3 toolbox (v3.0 RC2; Tournier et al., 2019). After denoising the dMRI scans (*dwidenoise* in MRtrix), eddy current and motion correction was conducted (*eddy openmp* in FSL, with second level modeling, interpolation for outlier frames, and default setup otherwise). Then, the skull was stripped from eddy-corrected dMRI scans (*bet* in FSL, fractional intensity threshold = 0.3) and a white-matter mask, obtained from segmented anatomical images (using SPM12 *Segmentation* algorithm, Ashburner and Friston, 2005), was mapped to the resolution of dMRI scans (*flirt* in FSL) and dilated by one voxel (*maskfilter* in MRtrix). Then, we estimated the single-bundle response function for spherical deconvolution based on the Tournier algorithm (Tournier et al., 2013) and computed the fibre orientation distribution function for every voxel with a constrained spherical deconvolution (CSD; Tournier et al., 2007). CSD-based deterministic fibre tracking (*SD Stream* in MRtrix; Tournier et al., 2012) was applied to reconstruct 10×10^6^ streamlines longer than 10 mm. This CSD-based fibre tracking approach has the advantage that it is capable of tracking trough regions of crossing fibres, thereby overcoming limitations of more traditional tensor based approaches. To correct biological inaccuracy related to inherent bias of the stream reconstruction algorithm, spherical-deconvolution informed filtering of tractograms was performed (SIFT; Smith et al., 2013). In this step, the reconstructed streamlines were reduced to a number of 1×10^6^ streamlines for each subject. The Brainnetome atlas (http://atlas.brainnetome.org) was warped from MNI-to subject-space and down-sampled to dMRI resolution using SPM12. Finally, a structural connectivity matrix 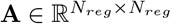 was reconstructed for every subject by counting the streamlines connecting each of the *N_reg_* = 234 regions in the Brainnetome atlas.

### 2.5. Minimum control energy to persist in a brain state

In this study, we used linear control theory for brain network analysis, an approach that uses principles from control and dynamical systems theory to investigate the impact that the brain’s structural topology may have on its functional dynamics (Kim and Bassett, 2019; Tang and Bassett, 2018; Lynn and Bassett, 2019). Under the assumption of a linear model of dynamics (Kim and Bassett, 2019), and control from all regions of the brain, we estimated the minimum control energy required to persist in specific brain states. For extended reviews on the control of brain network dynamics, we refer the interested reader to (Tang and Bassett, 2018; Lynn and Bassett, 2019). In the following paragraphs, we describe the linear dynamic model used here, and outline the mathematical basis for the computation of the minimum control energy based on this model, as well as the way in which brain states of interest were defined in order to estimate persistence control energy.

#### Dynamic model

In order to study how the white-matter anatomy of the brain constrains or facilitates state transitions, we modeled the brain as a continuous linear time-invariant dynamic system following Gu et al. (2017) and Betzel et al. (2016)

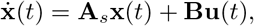

in which 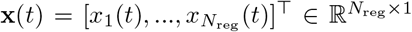 is the brain’s functional state at timepoint *t* given by the activity level *x_i_*(*t*) in each region *i*. The dynamic behavior of the brain is constrained by the stabilized structural white matter connectivity matrix **A**_*s*_, which is derived from the original structural connectivity matrix **A**, where each element *A_ij_* is the number of streamlines connecting regions *i* and *j*. In order to ensure stability of the dynamic system, all eigenvalues of **A**_*s*_ have to be below 0. Therefore, we stabilized the system by dividing the original structural connectivity matrix **A** by its largest eigenvalue *λ_max_* and subtract the identity matrix 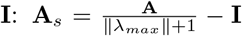 (Betzel et al., 2016). The diagonal matrix 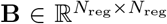 specifies the set of control nodes. Throughout this study, we assume that all regions of the brain can be controlled and therefore **B**= **I**. Finally, 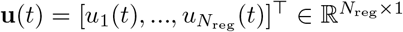 contains the control input signals *u_i_*(*t*) at region *i* and timepoint *t*.

Notably, the comparison of different graph models shows that there are marked differences in controllability across different types of networks, indicating that characteristics of brain network controllability are unique and potentially relevant for cognitive function (Wu-Yan et al., 2018).

#### Persistence control energy

In this study, we wished to investigate the structural control energy that is necessary to persist in a certain brain state **x** for a duration *T*. Throughout this study, we compute control energy for a control horizon of *T* = 1 (Betzel et al., 2016). The optimum control input **u**_*min*_ associated with minimum control energy can be found by solving (Antsaklis and Michel, 2005)

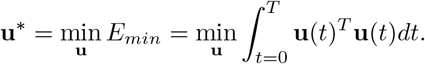

The analytical solution of this minimization problem is given by the minimum energy

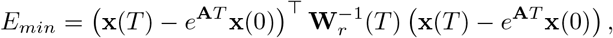

with the *reachability Gramian*

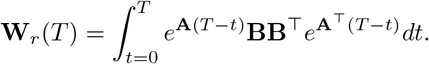

The optimal control input with this minimum energy can be explicitly computed for each timepoint *t*:

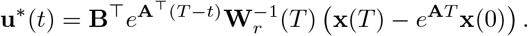

The required control energy at every brain region is given by 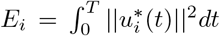 and the total control energy can be computed by summing over all regions 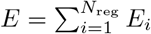.

Throughout this manuscript, we investigated persistence control energy, i.e., the minimum control energy in the case where initial and target states are identical: **x**(0) = **x**(*T*) = **x**. Then, minimum persistence control energy *E_P_* is defined by

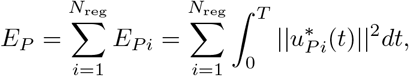

with

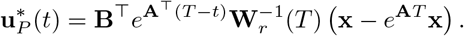

#### Definition of brain states

Persistence control energy *E_P_* as defined above was computed for every iCAP *k*. To this aim, brain state vectors **x**_*k*_(0) = **x**_*k*_(*T*) = **x**_*k*_ were obtained by computing for each of the 234 regions in the Brainnetome atlas the average normalized spatial map of every iCAP *k* across the voxels of the region (see subsection *fMRI processing*). In order to minimize noise susceptibility, we thresholded the brain states at a z-score of 1.5; in other words, regions with an average z-score *<* 1.5 were set to zero.

### 2.6 Statistical analysis

Statistical group comparison and brain-behavior analysis were conducted with an identical protocol as in Zöller et al. (2019).

#### Group comparisons of duration and persistence control energy

Two-sample t-tests were used to compare functional and structural measures between patients with 22q11DS and HCs, and permutation testing based on maximum T-statistics (Westfall and Young, 1993) using 5000 permutations was used to correct for multiple comparisons.

#### Partial least squares correlation

*multivariate relationship between brain measures and age.* We used partial least squares correlation (PLSC; Krishnan et al., 2011) to retrieve patterns of age-relationship in persistence control energy of all iCAPs. The steps of PLSC include

1. Computation of concatenated group-wise correlation matrices 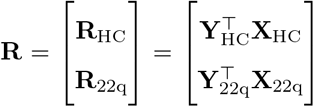 between age in **Y***∈* 1 *× N*_sub_ and persistence control energy in **X***∈ K × N*_sub_ of each brain state *k* = 1*, …, K* and subject *s* = 1*, …, N*_sub_. **Y**_HC_, **Y**_22q_, **X**_HC_, and **X**_22q_ are z-scored across subjects.
2. Singular value decomposition of **R**= **USV**^*T*^ to obtain a number of correlation components (also often called ‘latent variables’). Singular values on the diagonal of **S** indicate the explained correlation by each component. Every component is associated with one age weight per group (also often called ‘age saliences’) in **U** that indicate how strong the age relationship is present in each group, as well as with one brain weight per brain state (also often called ‘brain saliences’) in **V** that indicate how strongly each brain state contributes to the multivariate correlation between age and persistence control energy. These age weights and corresponding brain weights indicate the effect strength in each brain state. Because **X** and **Y**were z-scored in the first step, **R** corresponds to a correlation matrix, and as a consequence the age weights and brain weights can be interpreted similar to correlation values.
3. Permutation testing with 1000 permutations to test for significance of correlation components, where rows of **X** were randomly permuted, while leaving **Y** unchanged, in order to estimate the null distribution of singular values **S** under assumption of no significant correlation between **Y** and **X**.
4. Bootstrapping with 500 bootstrap samples, obtained through sampling with replacement the observations in **Y** and **X**, to evaluate the stability of age- and brain weights in a significant correlation component.

Altogether, PLSC allows to retrieve components that summarize the multivariate correlation between a number of behavioral measures and a number of brain measures. Each significant correlation component contains ‘behavior weights’ (here: age weights) and ‘brain weights’ (here: ‘persistence control energy weights’) that indicate, which behavior is most strongly correlated with which brain measure. For a more detailed description of PLSC, we refer the interested reader to (Krishnan et al., 2011; Zöller et al., 2017b,a, 2018).

#### Nuisance variable regression

Age and sex were included as nuisance regressors in group comparisons and only sex was used in age-PLSC analyses. Nuisance regressors were standardized within each group to avoid linear dependence with the effects of interest.

### 2.7. Relationship between brain state function and structure

In order to assess the relationship between resting-state fMRI activation measures and persistence control energy, we compute Pearson correlations between the two measures. First, we computed correlations across subjects for each group and each brain state, resulting in *K* = 17 correlation values per group. We then compared these 17 values between HCs and patients with 22q11DS using a paired t-test. Second, we computed correlations across the *K* = 17 brain states for each subject. We then compared the correlation values between HCs and patients with 22q11DS using a two-sample t-test.

## 3. Results

### 3.1. Spatial and temporal properties of resting-state functional states

Using iCAPs, we extracted 17 functional brain states from the resting-state fMRI scans. The optimal number of states was determined using consensus clustering (Monti et al., 2003). Extracted networks include well-known resting-state brain states, such as DMN, FPN, and salience network (SN) states (see supplementary figure S1). Patients with 22q11DS have significantly altered activation duration in 5 brain states, including both brain states with longer activation duration, and brain states with shorter activation duration (see supplemetary figure S2). For a detailed discussion of temporal properties and their relevance for clinical symptoms in 22q11DS, we refer the interested reader to (Zöller et al., 2019).

### 3.2. Persistence control energy in brain networks is lower than in random networks

We computed persistence control energy of the 17 brain states for every participant based on his or her individual structural connectivity matrix. In order to verify whether the persistence control energy values that we measure for human brain networks are different from random networks, we computed for every brain state the persistence control energy based on 100 random null networks that preserve the structural network topology (Rubinov and Sporns, 2010). For all states, the energy for these random networks was significantly higher than for the true brain networks (see supplementary figure S5), confirming previous findings by Cornblath et al., 2020 who also observed that brain networks are specifically wired to reduce control energy.

To furthermore test how specific the energy is to the particular data-driven resting-state brain states, we computed the persistence control energy for 100 brain states, in which regional brain state activation was randomly shuffled. For all states, the persistence control energy for randomly shuffled states was significantly higher than for the true brain state (see supplementary figure S6), indicating that the spatial distribution of functional brain states reduces persistence control energy given the brain’s structural topology.

### 3.3. Persistence control energy of functional brain states is altered in patients with 22q11DS

Aberrant structural connectivity in patients with 22q11DS leads to altered persistence control energy in four out of the 17 brain states (see figure 1). Persistence control energy was higher in patients with 22q11DS in brain states that involve more posterior and dorsal regions – primary visual 2 (PrimVIS2) and auditory / sensorimotor (AUD/SM). Reductions of persistence control energy on the other hand were present mainly in brain states including more anterior regions – dorsal anterior cingulate cortex /dorsolateral prefrontal cortex (dACC/dlPFC) and anterior DMN (aDMN).

**Figure 1:**
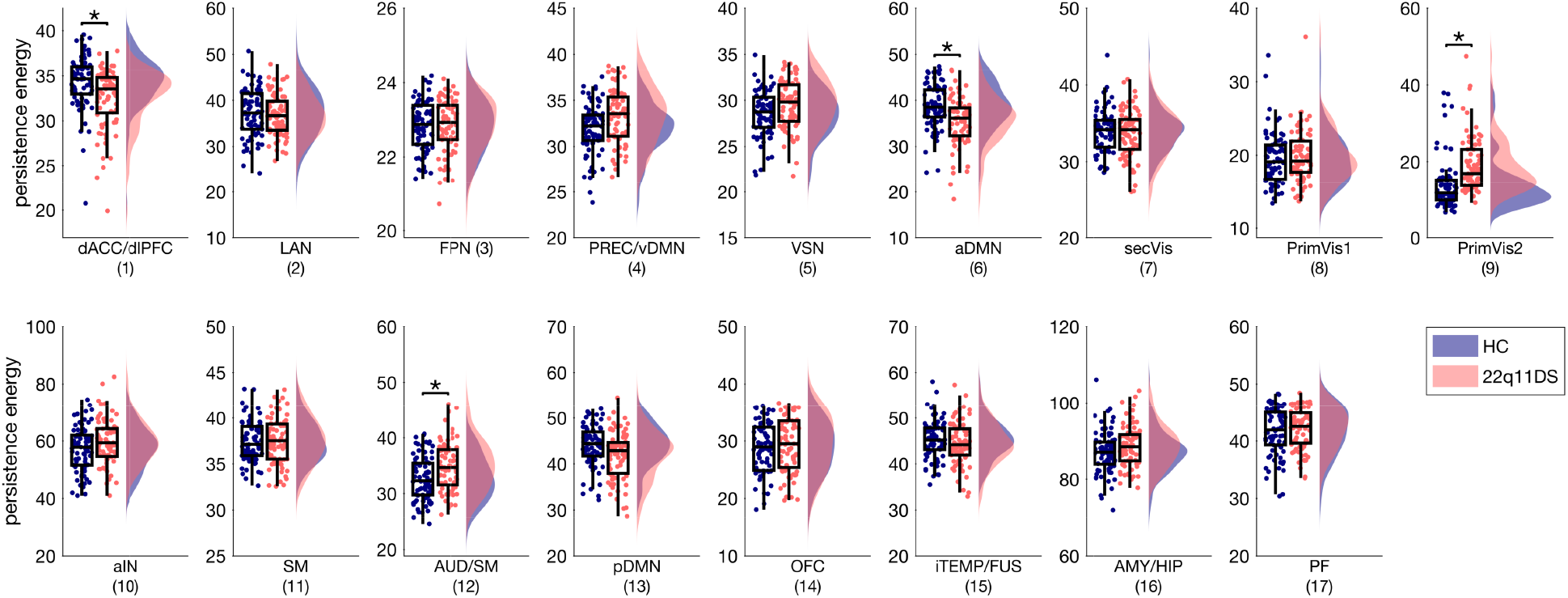
Group differences in persistence control energy of the 17 functional brain states in patients with 22q11DS compared to HCs. P-values are corrected for multiple comparisons based on permutation testing (Westfall and Young, 1993), age and sex were included as covariates. Significant group differences (p*<*0.05) are marked with an asterisk. Single-subject duration measures are included as scatterplots. dACC/dlPFC – dorsal anterior cingulate cortex / dorsolateral prefrontal cortex, LAN – language network, FPN – fronto-parietal network, PREC/vDMN – precuneus / ventralDMN, VSN – visuospatial network, aDMN – anterior DMN, SecVIS – secondary visual, PrimVIS1 – primary visual 1, PrimVIS2 – primary visual 2, aIN – anterior insula, SM – sensorimotor, AUD/SM – auditory / sensorimotor, pDMN – posterior DMN, OFC – orbitofrontal cortex, iTEMP/FUS – inferior temporal / fusiform, AMY/HIP – amygdala / hippocampus, PFC – prefrontal cortex.

### 3.4. Structural connectivity (weighted degree) alterations are inversely related to persistence control energy alterations

As we investigate persistence control energy allowing all brain regions to be controlled, and using a linear dynamic model, we expect persistence control energy to be closely related to the weighted degree of the brain regions in each brain state (Karrer et al., 2019). The specific relationship between structural connectivity and controllability depends on the model that is used (Karrer et al., 2019). Therefore, and to investigate this aspect in more depth, we computed the weighted sum of the regional weighted degree, multiplied by the spatial map of each brain state. Results of univariate t-tests comparing the weighted degree of brain states between groups (supplementary figures S3) show significant group differences between patients with 22q11DS and healthy control subjects in three brain states that all have also significantly altered persistence control energy (higher in dACC/dlPFC and aDMN, and lower in PrimVis2). These results indicate that reduced persistence control energy in patients with 22q11DS is caused by higher structural connectivity of the corresponding brain states, whereas higher persistence control energy in other brain states is caused by weaker structural connectivity of these states.

### 3.5. Persistence control energy decreases from childhood to adulthood

There is one significant correlation component (p*<*0.001) resulting from PLSC analysis testing for a relationship between persistence control energy and age (see figure 2). Persistence control energy is negatively correlated with age in 7 out of the 17 states, as reflected by negative iCAPs persistence control energy weights (figure 4B). This correlation with age is significant for both groups, but stronger in patients with 22q11DS than in HCs, as indicated by the stronger age weights in patients with 22q11DS than healthy control subjects (figure 4A). Brain states that show a stable relationship between persistence control energy and age include anterior and posterior DMN, anterior insula (aIN), sensory states (primary visual 1 (PrimVIS1) and sensorimotor (SM)), one emotional state (AMY/HIP), and the language state. The persistence control energy of states that are commonly associated with goal-directed behavior during tasks do not show any relationship with age (i.e., precuneus / ventral DMN (PREC/vDMN), which is sometimes called dorsal attention state, FPN, and visuospatial network (VSN))

**Figure 2:**
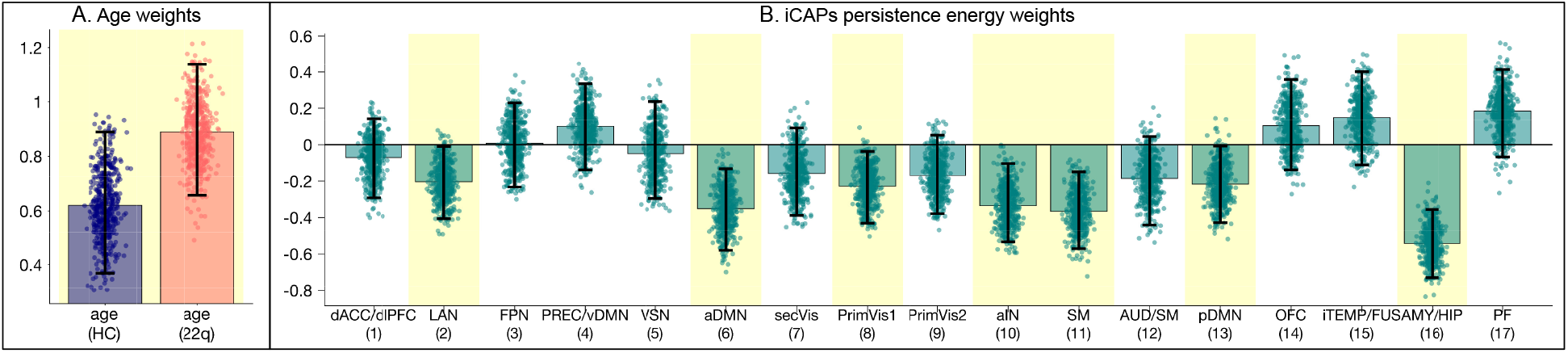
PLSC testing for a relationship between persistence control energy and age resulted in one significant correlation component (p*<*0.001). A) According to age weights indicating the correlation strength in each group, the age-relationship is stronger in patients with 22q11DS than in HCs. B) Persistence control energy weights show that there is a stable negative relationship with age in 7 out of the 17 brain states. Error bars indicate bootstrapping 95% confidence intervals; stable results were indicated by yellow background.

### 3.6. No correlation across subjects between activation duration and persistence control energy

In order to test whether there was a relationship between structural and functional measures across subjects, we computed correlations between resting-state activation duration and persistence control energy across subjects for each state. As group differences in persistence control energy (figure 1) were not present in the identical states as group differences in resting-state activation (supplementary figure S2), we expected no linear correspondence between the two measures. Indeed, there was no significant correlation between the two measures either in patients with 22q11DS or in HCs (see figure 3A). In other words, a subject who spent a long time in one brain state during the resting-state fMRI scan (measured through iCAPs activation duration), did not have a systematically higher or lower persistence control energy (based on the subject’s structural connectivity and a model of brain dynamics) of that brain state compared to other subjects.

**Figure 3:**
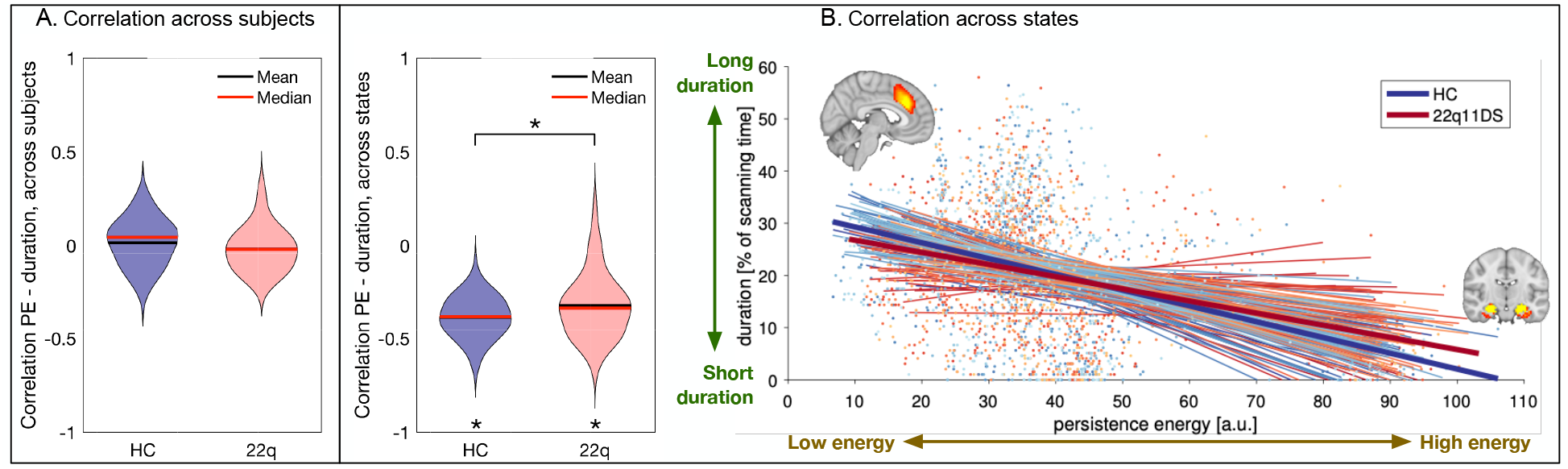
Correlation between persistence control energy and resting-state activation duration. A) Across subjects there is no significant correlation, either in patients with 22q11DS or in HCs. Violin plots show the distribution of correlations for all brain states. B) Across states there is a negative correlation between the two modalities: the higher the persistence control energy, the shorter the resting-state activation duration. This correlation is significantly stronger in HCs than in patients with 22q11DS. Left: Violin plots show the distribution of correlations for all subjects. Significant group differences (p*<*0.05) are marked with an asterisk. Right: every line corresponds to the fitted linear curve for each subject, thick lines show the average correlation of each group. PE – persistence control energy.

**Figure 4:**
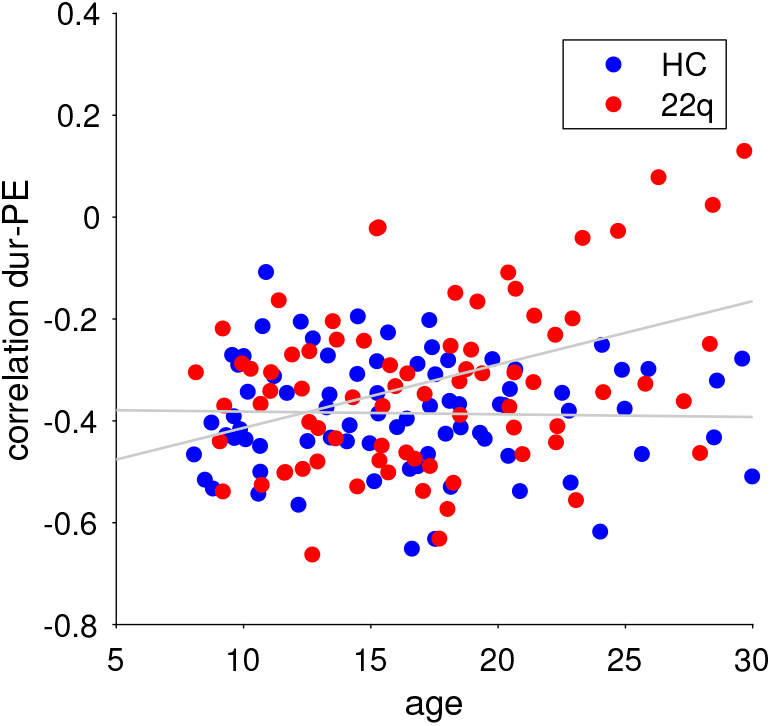
The negative correlation between persistence control energy and activation duration across brain states is constant over age in HCs (c=−0.03, p=0.937), but significantly increasing (i.e., weakening) with higher age in patients with 22q11DS (c=0.38, p=0.005). dur – activation duration; PE – persistence control energy.

### 3.7. Negative correlation across states between activation duration and persistence control energy

To test whether within each subject there was a relationship between temporal activation and persistence control energy, we computed across-states correlations for each subject (see figure 3B). There was indeed a negative correlation between activation duration and persistence control energy. In other words, all subjects tend to spend less time in brain states whose structural wiring leads to higher persistence control energy. This negative correlation was significantly stronger (lower) in HCs than in patients with 22q11DS (p=0.010). To test whether this correlation between structure and function changes over age, we tested for a relationship with age of these negative correlation values (see figure 4). In HCs the energy-activation correlation did not change over age (c=−0.03, p=0.937). In patients with 22q11DS, however, the correlation became significantly weaker (higher negative values) with increasing age (c=0.37, p=0.005). In a joint model, the group-by-age interaction was significant with p=0.002. In other words, while at a younger age, patients showed similar energy-activation relationship as HCs, older patients have a weaker negative energy-activation correlation, indicating that the observed weakening in correlation (see previous paragraph) is emerging over age.

There was no correlation with full-scale intelligence quotient (FSIQ; HCs: c=0.08,p=0.728; 22q11DS: c=−0.01, p=0.0.937), positive psychotic symptoms as measured by the sum of positive SIPS scores (22q11DS: c=−0.12, p=0.688) and negative psychotic symptoms as measured by the sum of negative SIPS scores (22q11DS: c=−0.37, p=0.130). Reported p-values were corrected for the six computed correlations using the FDR.

## 4. Discussion

Here we used dynamic models based on network control theory to investigate the impact of aberrant structural connectivity on the activation of functional brain states during rest. We investigated whether the relationship between structure and function is altered in patients with 22q11DS, a population at extremely high risk for schizophrenia. Our overall goal was to test whether such an alterations could give insights on possible mechanistic implications of aberrant brain structure on altered functional activation (Braun et al., 2018).

To address this question, we retrieved functional brain states and their activation duration from resting-state fMRI scans. Then, we used network control theory to predict the control energy that would be necessary to persist in these functional states based on structural connectivity measured in diffusion MRI (Braun et al., 2019; Cornblath et al., 2020). Finally, we investigated the relationship between the temporal activation during rest and this predicted persistence control energy, in order to estimate how the brain’s white matter anatomy may influence the dynamic behavior of functional brain states. As this is one of the first studies using network control theory to investigate a clinical population, our aim was to explore whether and how this approach could give insights into aberrant structure-function relationships in psychiatric disorders. We found aberrant persistence control energy of several brain states, mainly involving anterior or posterior medial connections. Further, persistence control energy decreased from childhood to adulthood both in patients with 22q11DS and HCs. Finally, when probing for a relationship between structural persistence control energy and functional resting-state activation, we found a negative correlation across brain states consistent with prior work (Cornblath et al., 2020), which was less pronounced in patients with 22q11DS. However, even though in some brain states both persistence control energy and functional activation were altered in patients with 22q11DS, we found no systematic relationship between persistence control energy of a brain state and its functional activation across subjects. In the following exposition, we will first discuss the alterations of persistence control energy and altered structure-function relationships in patients with 22q11DS, and then offer a tentative explanation for the absence of an across-subject relationship between control energy and functional activation.

### 4.1 The architecture of human structural brain networks and functional brain states support low persistence control energy

Overall, we confirmed previous findings comparing persistence control energy in human brain networks with those of randomized networks (Cornblath et al., 2020). For both patients with 22q11DS and healthy controls, persistence control energy was lower in real brain networks than in topology-preserving random networks. Also persistence control energy for the true data-driven brain states was lower than for spatially shuffled brain states. Together, these results indicate that both human structural white matter topology, and functional brain state architecture, support the engagement in brain states at low energetic cost.

### 4.2. Anterior-posterior and medial-lateral gradient of altered connectivity leads to aberrant brain control energy in 22q11DS

The first goal of this study was to investigate persistence control energy of data-driven resting-state functional brain states in patients with 22q11DS. We found that aberrant structural wiring leads to a pattern of altered control properties with some brain states requiring higher persistence control energy and others requiring lower persistence control energy. Persistence control energy is mainly reduced in frontal brain states (aDMN, dACC/dlPFC) and increased in occipital (PrimVIS2) and lateral parietal (AUD/SM) brain states. These alterations were inversely related to altered weighted degree, for which frontal brain states (aDMN, dACC/dlPFC) showed increases, whereas the occipital PrimVIS2 brain state showed reductions. This pattern of findings confirms prior reports of aberrant anterior-posterior and medial-lateral white-matter connectivity in patients with 22q11DS, which was found to relate to positive and negative psychotic symptoms (Scariati et al., 2016; Gothelf et al., 2011; Váša et al., 2016). Our study expands upon these prior studies by probing the impact of this aberrant wiring on the dynamic behavior of the brain in terms of the predicted persistence control energy that is needed to engage in these brain states. Profound reductions of persistence control energy were present in DMN and cingulo-frontal SN, which are two brain systems that are known to play a central role in higher order cognition (Menon, 2011). Previous reports provide evidence that the structural and functional connectivity of these brain states is altered in patients with 22q11DS (Padula et al., 2015, 2017b; Schreiner et al., 2014). In particular the dACC, which is a central node of the SN, has been found to be affected in 22q11DS using different neuroimaging modalities and alterations of this brain region have been suggested as a biomarker for psychosis in the disorder (Padula et al., 2018). Lower persistence control energy in these brain states in patients with 22q11DS may seem counterintuitive, as the healthy brain is assumed to decrement energy. Speculatively, the effect could be related to compensatory effects in less symptomatic patients. However, patients with 22q11DS do not spend more time in these energetically more advantageous states, so even though these brain states should be more easily accessible, they do not make use of this advantage. Further research on the relationship between persistence control energy and different clinical profiles of individuals with 22q11DS would be promising to investigate this effect in more detail.

### 4.3. The brain gets energetically more efficient from childhood to adulthood

Aside from alterations in 22q11DS, we found that persistence control energy of many brain states (7 out of 17) is negatively correlated with age, both in patients with 22q11DS and in HCs. This finding suggests that with increasing age, the brain gets more efficiently wired to reduce the control energy required for its functional activation. In line with these findings, Tang et al. found that both average and modal controllability (which measure the general ability to steer the brain towards easy-to-reach or difficult-to-reach brain states) increase over age in a similar age range, which suggests an increasingly efficient wiring that at the same time allows a higher diversity of brain dynamics (Tang et al., 2017). Importantly, while there are small variations in the distributions of controllability across the brain in males and females, the trend for increasing controllability with age is equally strong in both sexes (Cornblath et al., 2018). Further, Cui et al. (2020) found that control energy of atlas-based brain states calculated with a comparable approach to ours, was also decreasing from childhood to adulthood in most brain states (Cui et al., 2020). This developmental trajectory of structural brain architecture was preserved in patients, suggesting that while they present with absolute alterations of controllability properties, their overall development seems to be largely intact.

### 4.4. Dynamic inefficiency in patients with 22q11DS, which becomes increasingly marked with age

When investigating the relationship between functional activation and structural persistence control energy, we found that the brain activates in a highly efficient way, spending less time in brain states that are energetically more demanding, consistent with prior evidence in a completely different cohort (Cornblath et al., 2020). In patients with 22q11DS, this relationship was significantly weaker than in HCs, which suggests that aside from the pure alteration in structure and function, the relationship between the two is also altered. In particular, patients use their brains in a less efficient way, spending more time in energetically demanding states than HCs. Additionally, this dynamic inefficiency in patients with 22q11DS became increasingly marked with greater age. Possibly, the patients’ inefficient use of their brain may express in the more severe symptomatology characteristic of older patients. Divergent trajectories during adolescence have for instance been reported for executive functions (Maeder et al., 2016). However, here we did not find a significant correlation between our measure of dynamic efficiency and psychotic symptoms or IQ. The absence of such behavioral relationships could indicate that this measure of dynamic efficiency represents a general property of human brain structure-function relationship, which is preserved even in the presence of clinical symptoms. Contrarily to structural connectivity and functional brain state activation separately, their relationship as measured here, does not seem to be a measure sensitive to the symptomatology of schizophrenia. However, it should be noted that individuals with 22q11DS share a common and broad genetic vulnerability towards multiple forms of psychopathology. In this perspective, lack of clinical correlations could imply that a dynamic inefficiency of resting-state networks represents a trait marker of vulnerability to psychopathology rather than a state marker related to the presence of symptoms at the time of scanning. Nevertheless, it is noteworthy that here we present an initial test in a sample of limited size, and only testing for global summary measures of psychosis. Future studies with larger sample sizes, and targeting more specific measures of particular clinical symptoms and executive functions, would be required to draw a final conclusion on this.

### 4.5. Control energy and activation alterations manifest in similar brain states, but are not systematically correlated across subjects

While we were able to detect a relationship between functional activation and structural persistence control energy across states (pointing towards the inherent efficiency of the human brain as discussed above), we did not observe a relationship across subjects. In other words, subjects with less efficiently wired brain states (higher persistence control energy) did not systematically spend more or less time in these brain states compared to other subjects. This can also be seen when comparing the alterations that we find in persistence control energy with results from our previous study, where we investigated functional properties in the identical subjects by also using the iCAPs approach to assess temporally overlapping brain state activation (Zöller et al., 2019). As indicated by the absence of across-subject correlations (figure 3A), the direction of alterations of resting-state activation duration obtained in that study was not linearly related to structural persistence control energy alterations found in the present study. This suggests that the measure of iCAPs activation duration, and persistence control energy represent different types of brain alterations in these patients. However, there were overlapping results in some of the brain states, which indicates that in these particular brain states, both structural and functional properties play a role in the 22q11DS symptomatology.

In particular, in this previous study we found a relationship between altered resting-state activation in DMN and cingulo-frontal SN and higher levels of anxiety (anterior DMN), and positive psychotic symptoms (cingulo-frontal SN) (Zöller et al., 2019). Furthermore, and in line with the literature on the role of the amygdala and hippocampus in anxiety (Etkin and Wager, 2007), we found that higher functional activation and aberrant functional coupling of the amygdala and hippocampus brain state tracks with higher levels of anxiety. In the present study, we confirm that DMN and cingulo-frontal SN are also affected in terms of structural connectivity measured through diffusion MRI, and that this alteration in structural wiring leads to aberrant controllability properties in patients with 22q11DS. Future studies should examine whether these alterations are also linked to higher levels of anxiety and the risk for developing psychosis as suggested by our previous functional and anatomical findings (Zöller et al., 2019; Mancini et al., 2019).

Importantly, brain states do not act in isolation, but interact among themselves. For instance, knowing about the interaction between the DMN and the amygdala and hippocampus that we discovered in the same dataset (Zöller et al., 2019), one could imagine that indeed alterations of controllability in the DMN may lead to increased activity of the amygdala and hippocampus. Testing for such cross-network relationships may be even more interesting, and our methological approach offers a valuable framework to explore this complexity in future research.

### 4.6. Significance and implication for the pathophysiology in schizophrenia

Altogether, the results presented here show that network control theory, and in particular persistence control energy, can provide insights into global structure-function relationship of the human brain. Through the assumption of a dynamic model of brain function, it allowed us to go beyond a pure description of structure-function relationship in specific brain states, and provide a prediction of the effect that altered structural wiring has on the brain’s dynamic activation in terms of energy cost. Our results suggest that a negative relationship between functional activation and energy cost across brain states is global property of the brain’s function in every individual. The overall reduction of this relationship in a population at risk for schizophrenia suggests that psychiatric disease affects the brain in a way that makes it’s functional activation more energetically costly. However, this relationship did not relate to clinical variables, which could mean that this property represents a trait marker of vulnerability to psychopathology rather than a state marker related to the presence of symptoms at the time of scanning. A recent study investigating network control energy in patients with schizophrenia using a very similar approach to the one used here found increased control energy in a task-related working-memory brain state in patients with schizophrenia (Braun et al., 2019). These results also point towards an energetically more demanding structural wiring of the brain in patients with schizophrenia. Together with the dysfunctional activation of structurally affected brain states that we find here, these results provide an interesting starting point for future analyses of brain structure and function in schizophrenia, and psychiatric diseases in general. Especially an investigation of larger samples, and possibly also task-fMRI data, would be promising to confirm these findings, and get further insights into this aberrant relationship between functional activation and structural architecture.

### 4.7. Methodological considerations and limitations

#### Placement among existing state-of-the-art control strategies

The approaches using control theory to analyze brain networks can be differentiated based on the control strategies they use. Initial studies on controllability of brain networks examined properties like average and modal controllability, which assess the ease of a local or distant state transition while ignoring the nature of either the initial or target states (e.g., Gu et al., 2015; Tang et al., 2017; Jeganathan et al., 2018). Studies on specific state trajectories, on the other hand, stipulate the initial and target states and assess the energy necessary for that specific trajectory (Gu et al., 2017; Betzel et al., 2016; Cui et al., 2020). In this case, states could be defined either by activating all regions in a cognitive system defined from the literature (Gu et al., 2017; Betzel et al., 2016; Cui et al., 2020), or by choosing a data-derived brain state (Cornblath et al., 2020; Braun et al., 2019). Similar to the latter two studies (Cornblath et al., 2020; Braun et al., 2019), here we investigated transitions between specific states, and we defined the brain states in a data-driven way from fMRI data acquired from the same subjects. Using this approach, we found profound differences in control energy of multiple states in patients with 22q11DS. These findings underline the potential of data-driven brain states to detect relevant subject-specific alterations, which is promising for future studies involving other clinical populations.

#### Variance of structural connectivity across subjects and across regions has different scales

In the present study, structural connectivity was measured in terms of connection density; that is, with a fixed number of reconstructed streamlines. As a result, the variability of connectivity across regions is relatively high compared to the variability across subjects. Therefore, this approach supports a careful investigation into *relative* changes in connectivity, but it is less powerful in tracking *absolute* changes in connectivity. In our results, this effect can be observed in figure 1, where the differences in energy from one state to another are much larger than the differences in persistence control energy across subjects for one single brain state. Significant differences between subjects do exist, however, they are small with respect to the differences between brain states. This effect may be a possible reason why we detected correlations with functional activation across brain states, but not across subjects.

#### Linear models of brain dynamics

For simplicity, here we chose to use a linear model of dynamics on the structural connectivity graph (Kim and Bassett, 2019; Honey et al., 2009; Gu et al., 2015) to calculate minimum control energy (Betzel et al., 2016). Even though this model is the most widely used approach for network control theory in neuroscience, it may be overly simplistic. Incorporating models of non-linear dynamics could prove useful in the future as they could potentially improve the estimation of more realistic control energy (Kim and Bassett, 2019). It is possible that an estimate based on more biologically plausible dynamic models would allow us to detect more subtle relationships between controllability and functional activation.

#### Relevance for other imaging modalities

While the results presented here were obtained for fMRI brain states and thus specific to resting-state fMRI, the network control framework is not specific to fMRI and could also be applied to other types of data. In this regard, the framework has already used to study brain states derived from electrocorticography (Stiso et al., 2019), and could also be imagined for electroencephalography data.

#### Assessing structural connectivity through diffusion MRI

As the clinical dataset studied here was recorded as part of a large, longitudinal study for which the first scans were recorded more than 10 years ago, the diffusion MRI sequence constitutes a limiting factor. While motion may have had an influence on our result (Baum et al., 2018), only one b0 image was recorded in our diffusion MRI sequence, which made it impossible to estimate motion over the scanning duration. Therefore, motion may represent a confounding factor in our analysis that could not be quantified. Of note, another study investigating control energy who did control for motion also found decreasing control energy from childhood to adulthood (Cui et al., 2020), which suggests that in-scanner head motion was likely not a major confound in our study. To overcome limitations regarding streamline reconstruction to assess structural connectivity, we used tractography based on fibre orientation distribution and constrained spherical deconvolution (Tournier et al., 2013, 2007, 2012), in combination with SIFT to increase biological accuracy (Smith et al., 2013). Therefore, we believe that our streamline measure is a useful approach to estimate structural connectivity from the diffusion-weighted images in this dataset. However, there remain the general limitations related to diffusion imaging for structural connectivity reconstruction, and our results should be replicated in future studies using more advanced diffusion image acquisition protocols that allow for a better consideration of confounding factors.

## 5. Conclusion

In this study, we investigated the control energy of functional brain states in patients with 22q11DS. This is the first study investigating the impact of aberrant structural connectivity on brain dynamics using control energy in 22q11DS. We found that altered connectivity in patients with 22q11DS leads to reduced energy impact for engaging frontal brain states, whereas more occipital and parietal brain states were energetically more demanding for patients with 22q11DS than HCs. Further, in a comparison of structural persistence control energy with resting-state fMRI activation, we found that the brain functions in an efficient way by engaging less in energetically demanding brain states. In patients with 22q11DS the anticorrelation between activation and control energy is weaker than in controls, suggesting a dynamic inefficiency of brain function in these patients. In summary, we contribute one of the first studies investigating a direct link between control energy and functional activation during rest and provide promising insights for a better understanding of brain alterations in 22q11DS.

## Acknowledgements

We are grateful to the subjects who participated in our study and thank Sarah Menghetti, Léa Chambaz, Virginie Pouillard and Dr. Maude Schneider for their involvement with the participants. We would also like to acknowledge Prof. François Lazeyras and the CIBM group for their support during data collection.

This work was supported by the Swiss National Science Foundation (SNSF) under Grants 32473B 121966, 234730 144260 and 145250 to SE, and Grant 163859 to MS, and by the National Center of Competence in Research (NCCR) “SYNAPSY - The Synaptic Bases of Mental Diseases”, SNSF, under Grants 51AU40-125759, 51NF40-158776 and 51NF40-185897.

DSB acknowledges support from the John D. and Catherine T. MacArthur Foundation, the Alfred P. Sloan Foundation, the ISI Foundation, the Paul Allen Foundation, the Army Research Laboratory (W911NF-10-2-0022), the Army Research Office (Bassett-W911NF-14-1-0679, DCIST-W911NF-17-2-0181, W911NF-16-1-0474), the National Institute of Mental Health (2-R01-DC-009209-11, R01-MH112847, R01-MH107235, R21-M MH-106799), the National Institute of Child Health and Human Development (1R01HD086888-01), the National Institute of Neurological Disorders and Stroke (R01 NS099348), and the National Science Foundation (BCS-1441502, BCS-1430087, NSF PHY-1554488 and BCS-1631550). The content is solely the responsibility of the authors and does not necessarily represent the official views of any of the funding agencies.

## Data availability statement

The data that support the findings of this study are available on request from the corresponding author.

The data are not publicly available due to privacy or ethical restrictions.

## Diversity Statement

Recent work in several fields of science has identified a bias in citation practices such that papers from women and other minorities are under-cited relative to the number of such papers in the field (Mitchell et al., 2013; Dion et al., 2018; Caplar et al., 2017; Maliniak et al., 2013; Dworkin et al., 2020). Here we sought to proactively consider choosing references that reflect the diversity of the field in thought, form of contribution, gender, and other factors. We obtained predicted gender of the first and last author of each reference by using databases that store the probability of a name being carried by a woman (Dworkin et al., 2020; Zhou et al., 2020). The gender of authors whose names were classified as ‘unknown’ were found through online research. By this measure (and excluding self-citations to the first and last authors of our current paper), our references contain 12.5% woman(first)/woman(last), 18.2% man/woman, 22.7% woman/man, 46.6% man/man. This method is limited in that a) names, pronouns, and social media profiles used to construct the databases may not, in every case, be indicative of gender identity and b) it cannot account for intersex, non-binary, or transgender people. We look forward to future work that could help us to better understand how to support equitable practices in science.

