## supplementary material for "Structural control energy of resting-state functional brain states reveals inefficient brain dynamics in psychosis vulnerability"

**Contents**

**Supplementary tables** **2**

**Supplementary figures** **3**

**Supplementary references** **7**

*Corresponding author

*July 9, 2020*

### Supplementary tables

Table S1: Number of scans recorded before and after a scanner update. There was no significant group-by-scanner interaction.

| **image modality** | **Siemens Trio scanner (22q11DS/HCs)** | **Siemens Prisma scanner (22q11DS/HCs)** | **p-value ($\chi^2$)** |
|---|---|---|---|
| fMRI | 42/54 | 36/31 | 0.209 |
| dMRI | 41/49 | 36/29 | 0.227 |

### Supplementary figures

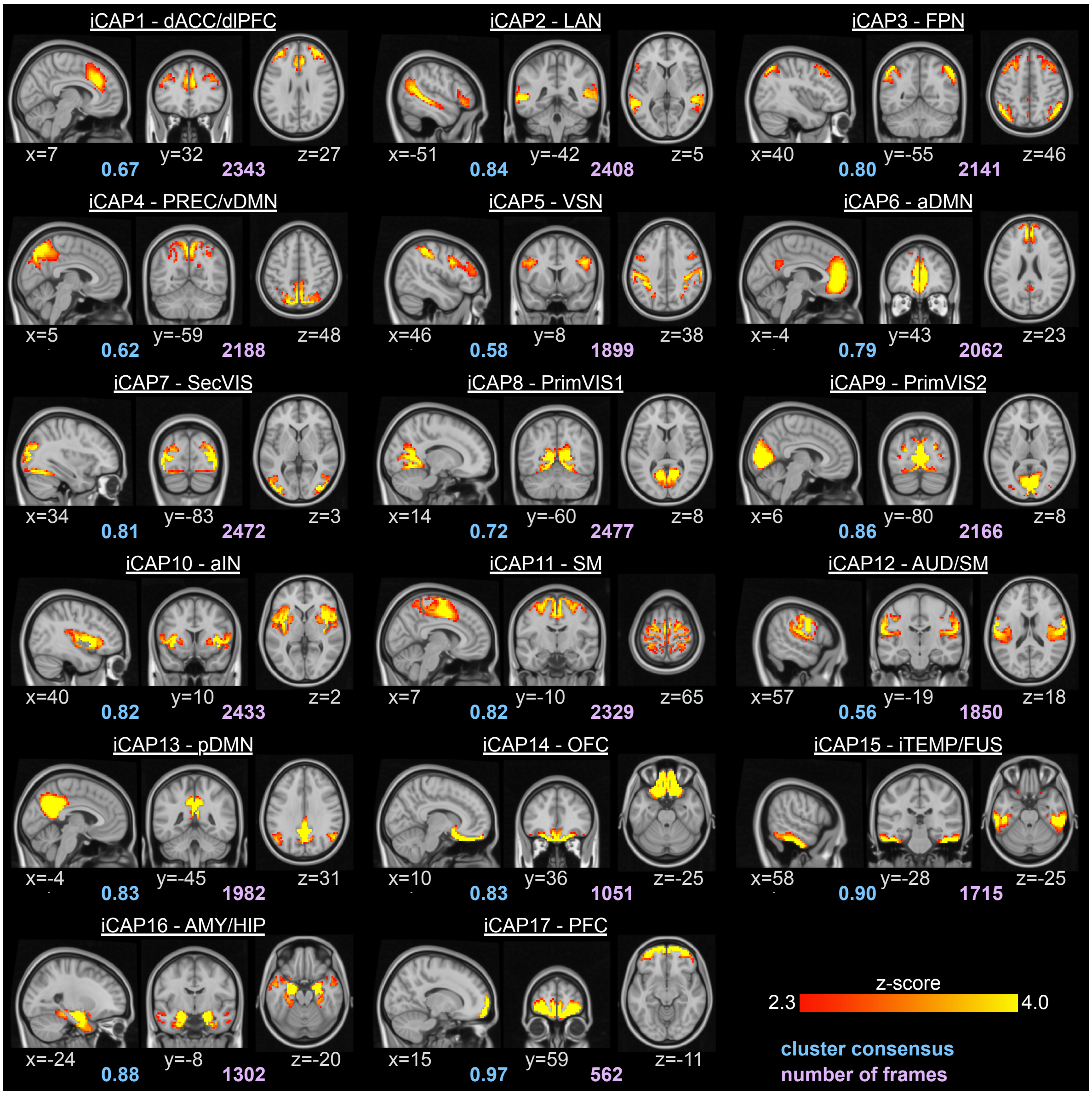


Figure S1: Spatial patterns of the 17 iCAPs retrieved from all subjects, including both HCs and patients with 22q11DS. The locations denote displayed slices in MNI coordinates. Blue values denote the average consensus of each cluster, purple values indicate the total number of innovation frames that were assigned to this cluster. Maps are identical to the ones in (Zöller et al., 2019), sorted according to the activation duration in HCs. dACC/dlPFC – dorsal anterior cingulate cortex / dorsolateral prefrontal cortex, LAN – language network, FPN – fronto-parietal network, PREC/vDMN – precuneus/ventral DMN, VSN – visuospatial network, aDMN – anterior DMN, SecVIS – secondary visual, PrimVIS1 – primary visual 1, PrimVIS2 – primary visual 2, aIN – anterior insula, SM – sensorimotor, AUD/SM – auditory/sensorimotor, pDMN – posterior DMN, OFC – orbitofrontal cortex, iTEMP/FUS –inferior temporal/fusiform, AMY/HIP – amygdala/hippocampus, PFC – prefrontal cortex.

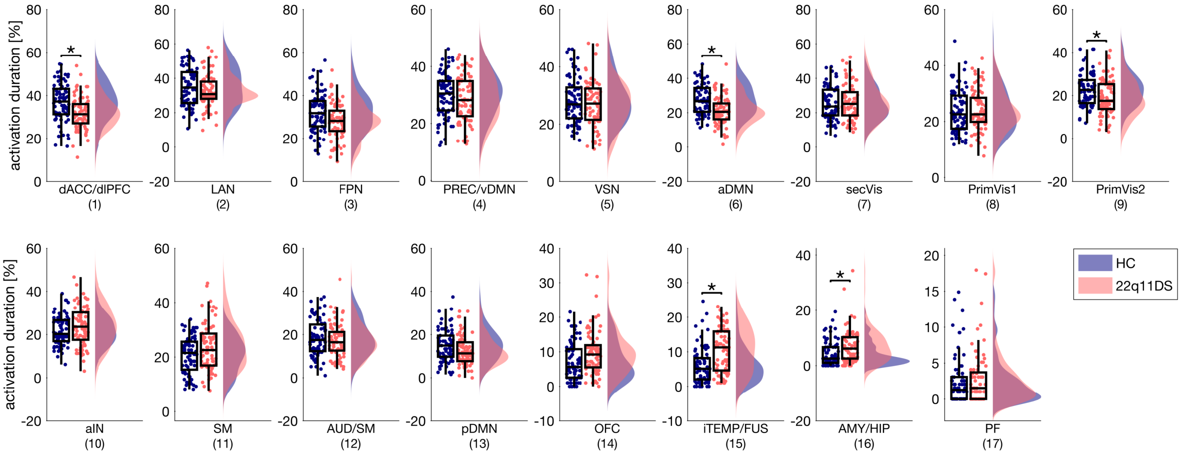


Figure S2: Statistics of total temporal duration for each iCAP. P-values are corrected for multiple comparisons based on permutation testing (Westfall and Young, 1993), and both age and sex were included as covariates. Significant group differences ($p<0.05$) were marked with an asterisk. Scatterplots represent the single-subject duration measures. Results are identical to those in (Zöller et al., 2019), but sorted according to the activation duration in HCs, and with alternative correction for multiple comparisons.

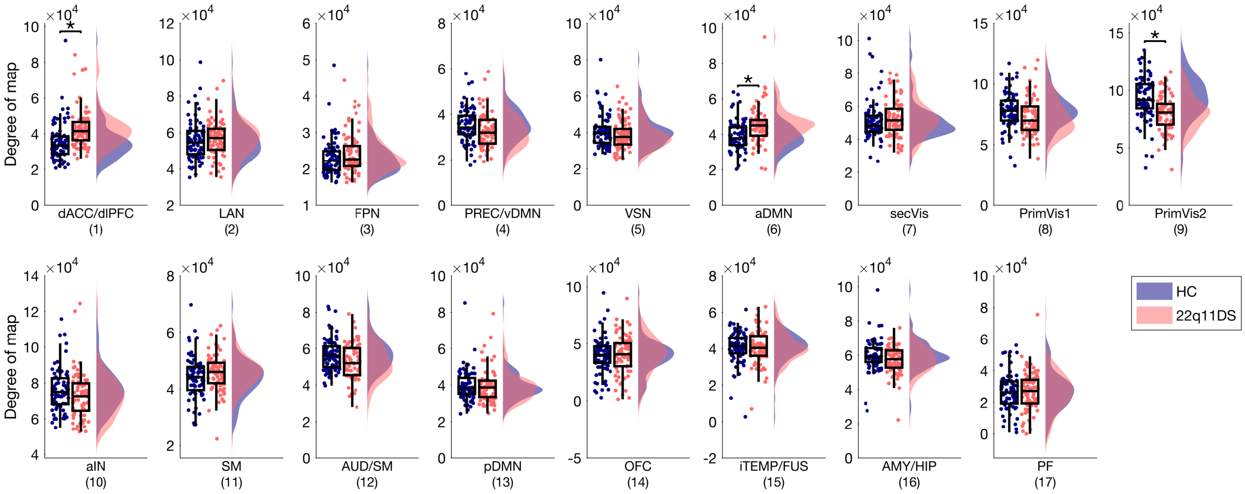


Figure S3: Group differences in degree (total streamline count per region) of the 17 functional brain states in patients with 22q11DS compared to HCs. Degree of each brain state was calculated as the sum of weighted degree per region, multiplied by the iCAP's regional map ($d_{iCAP} = \sum_{n=1}^{N_{reg}} d_n * z_{iCAP,n}$, with $N_{reg}$ the number of brain regions, $d_n$ the degree of the region, and $z_{iCAP,n}$ the value of the iCAP map in that region). P-values are corrected for multiple comparisons based on permutation testing (Westfall and Young, 1993), age and sex were included as covariates. Significant group differences ($p<0.05$) are marked with an asterisk. The results show that group differences in degree are significant in dACC/dlPFC, aDMN, and PrimVis2. All three of these brain states have also significantly altered persistence control energy (see figure 1 of the main manuscript), which is expected, due to the close relationship between persistence energy and weighted degree in whole-brain control problems (Karrer et al., 2019).

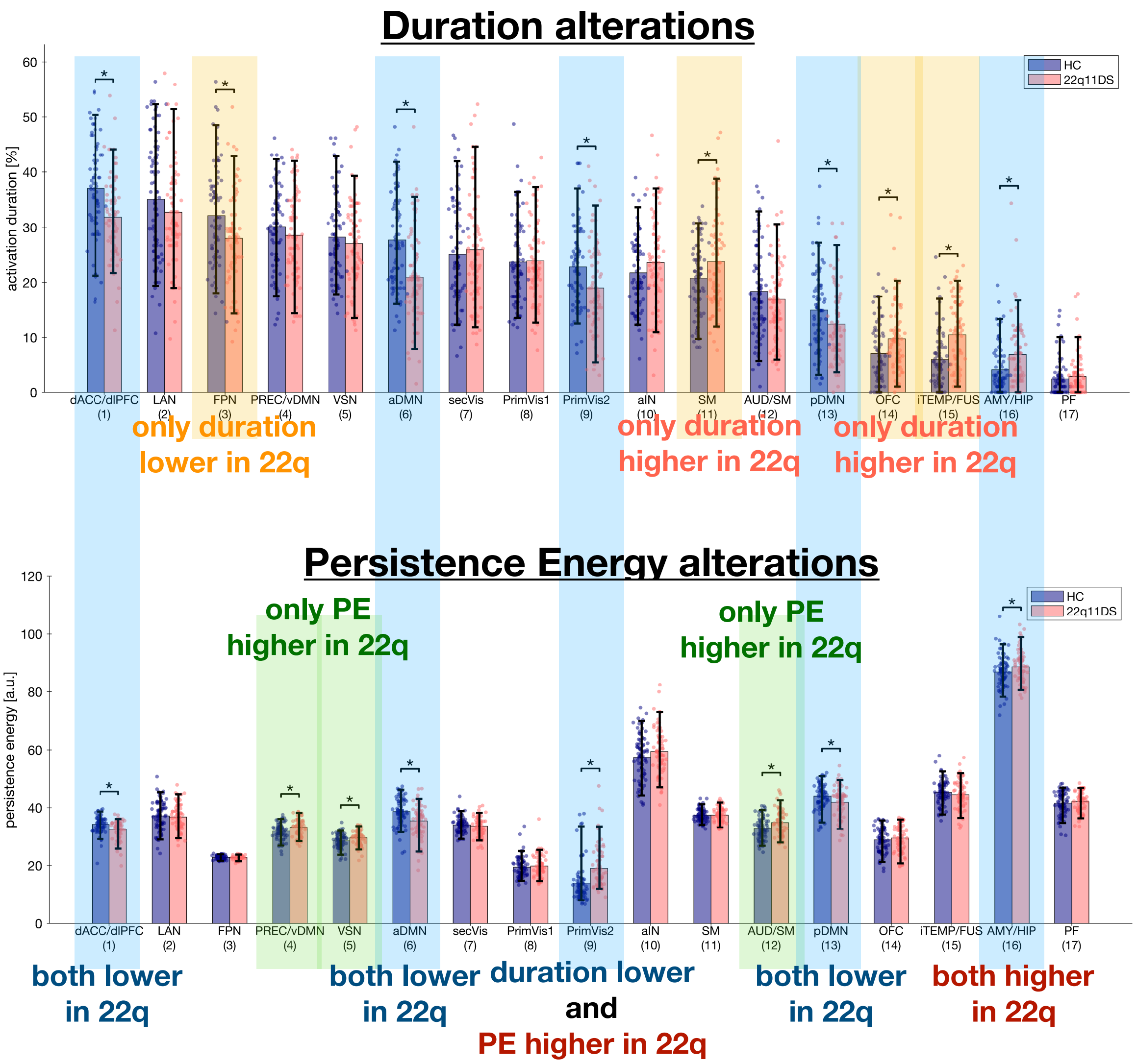


Figure S4: Comparison of alterations in resting-state activation duration and structural persistence control energy. While there were widespread alterations in both modalities, there was no clear pattern of common alterations.

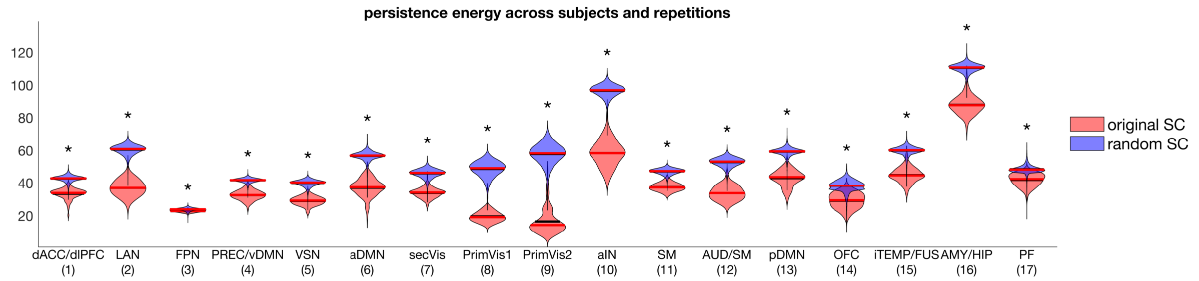


Figure S5: For each subject, persistence energy was computed for 100 random null models that preserve the subject's structural network topology (Rubinov and Sporns, 2010). The blue violin plots show the distribution of persistence energy values across the 100 repetitions and 155 subjects. The red violin plots show the original persistence energy of the 155 subjects.

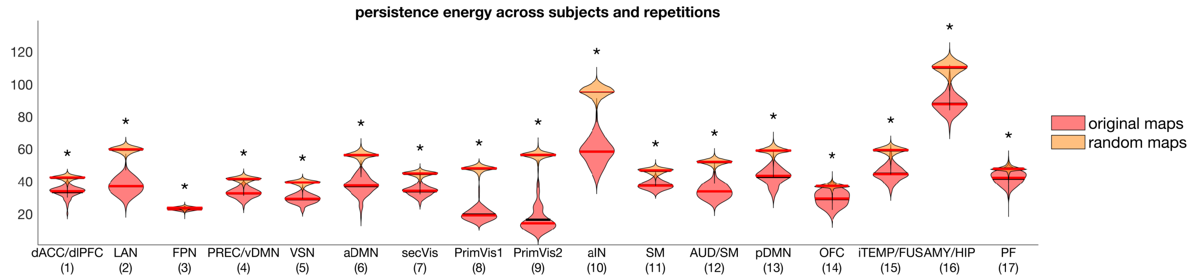


Figure S6: For each subject, persistence energy was computed pertaining the subject's structural connectivity, and 100 times randomly shuffling the iCAPs maps (random permutations of brain regions). The orange violin plots show the distribution of persistence energy values across the 100 repetitions and 155 subjects. The red violin plots show the original persistence energy of the 155 subjects.
